# AMES: Automated evaluation of sarcomere structures in cardiomyocytes

**DOI:** 10.1101/2021.08.06.455455

**Authors:** Maximilian Hillemanns, Heiko Lemcke, Robert David, Thomas Martinetz, Markus Wolfien, Olaf Wolkenhauer

## Abstract

**Background:** Arrhythmias are severe cardiac diseases and lethal if untreated. To serve as an in vitro drug testing option for anti-arrhythmic agents, cardiomyocytes are being generated in vitro from induced pluripotent stem cells (iPSCs). Unfortunately, these generated cardiomyocytes resemble fetal cardiac tissue rather than adult cardiomyocytes. An automated tool for an unbiased evaluation of cardiomyocytes would highly facilitate the establishment of new differentiation protocols to increase cellular maturity.

**Results:** In this work, a novel deep learning-based approach for this task is presented and evaluated. Different convolutional neural networks (CNNs) including 2D and 3D models were trained on fluorescence images of human iPSC-derived cardiomyocytes, which were rated based on their sarcomere content (sarcomerisation) and the orientation of sarcomere filaments (directionality) beforehand by a domain expert. The CNNs were trained to perform classifications on sarcomerisation, directionality ratings, and cell source, including primary adult and differentiated cardiomyocytes. The best accuracies are reached by a 3D model with a classification accuracy of about 90 % for sarcomerisation classification, 63 % for directionality classification, and 80 % for cell source classification. The trained models were additionally evaluated using two explanatory algorithms, IGrad and Grad-CAM. The heatmaps computed by those explainability algorithms show that the important regions in the image occur inside the cell and at the cellular borders for the classifier, and, therefore, validate the calculated regions.

**Conclusion:** In summary, we showed that cellular fluorescence images can be analyzed with CNNs and subsequently used to predict different states of sarcomere maturation. Our developed prediction tool AMES (https://github.com/maxhillemanns/AMES) can be used to make trustworthy predictions on the quality of a cardiomyocyte, which ultimately facilitates the optimized generation of cardiomyocytes from iPSCs and improves the quality control in an automated, unbiased manner. The applied workflow of testing different CNN models, adjusting parameters, and using a variety of explanatory algorithms can be easily transferred to further image based quality control, stratification, or analysis setups.

## Background

Differentiation, generation, and maturation of cardiomyocytes and pacemaker cells Cardiomyocytes, the muscle cells of the heart, can be generated by various methods in wet lab conditions: i) the differentiation of adipose tissue-derived mesenchymal stem cells [1], ii) the differentiation of murine or human embryonic stem cells [2, 3], and iii) the reprogramming of somatic cells [4], especially induced PSCs (iPSCs). It is also possible to extract adult cardiomyocytes from murine hearts for comparative analyses. These programming methods produce a cardiomyocyte aggregate, where some cells possess so-called pacemaker abilities and some do not. In recent approaches, the amount of non-pacemaker cells in this aggregate is still at around 20 % [5, 6]. Moreover, the electrophysiological properties of these cells resemble fetal cardiac tissue instead of adult cardiomyocytes [7]. The maturation level of cardiomyocytes may be critical for drug development, as immature cardiomyocytes are far more sensitive to potassium-channel blockers [8].

In general, fully developed cardiomyocytes possess a well aligned and highly organized sarcomere network [9]. Longer sarcomere structures correlate to an improved cardiac mechanical function [10]. Likewise, the mechanical, as well as the electrical function, are also dependent on the orientation of myofibrils and subsequently the sarcomere structures in a cell [11]. This orientation depends on the cell shape and the principal stress directions in the cell [12].

In order to validate and evaluate cardiomyocyte generation protocols, scientists need to examine the maturity of these cell aggregates. Hence, the need for an easily applicable method to distinguish between different maturation states is apparent [13].

### Cellular image analysis with deep learning

One possible approach to distinguish between different maturation states of cardiomyocytes can be the analysis of cellular images with a variety of deep learning (DL) applications. The main approaches commonly used refer to image segmentation (partitioning an image into meaningful parts or objects), object tracking (identifying and following an object through a time series), augmented microscopy (extraction of latent information from biological images), and, finally, image classification. Deep learning image classification has been used on a variety of different cells and tasks, like identifying changes in cell state [14], sorting cells into different phenotypes [15, 16, 17], and distinguishing between differentiated and undifferentiated cells on bright-field images [18]. By using DL, it is also possible to extract feature vectors from cellular images in order to cluster these vectors and gain insight on morphological patterns [19]. In comparison to classical machine learning (ML) approaches like support vector machines (SVMs) [20] or logistic regression [21], DL has shown to be more efficient at cell analysis tasks. In many cases, ML or DL is only applied after features were extracted from the images [20, 22]. In this study, DL was used directly on the images.

Transfer learning is also commonly used in biological image analysis due to the lack of available training data. In transfer learning, a neural network pretrained on another data set is applied to a new data set and the weights are retrained. Convolution Neural Networks (CNNs) are neural networks, which are able to extract features or patterns from images themselves, without any need for sophisticated preprocessing. CNNs combine this extraction part with the ability to make classification based upon these extractions. They were introduced by Bengio and Lecun in 1997 [23]. In cellular image analysis, CNNs are mainly used for image segmentation and not image classification [24, 25]. For image classification, fully connected layers are transferred after the convolutional layers to translate these features into a label. The labels that will be used here are different maturation degrees of sarcomere structures that have been introduced by a biological domain expert.

### Explainability analysis of image analysis models

As CNNs modulate highly nonlinear functions, they are too complex to allow for straightforward interpretability. This is referred to as the black box problem. A classifier may produce good classification results, but its inner workings and reasoning are unattainable [26]. The question ”Why does this model decide the way it does?” plays an increasingly important role, especially in the life sciences. In the last few years, a lot of algorithms have been developed to lift the lid of the black box. They can be sorted by two main criteria: local vs. global explanation and model-specific vs. model-independent. Local explanations are computed with the model, a data point and an output. In most cases, this is the predicted output of the model, although a different label can be used to find weaknesses in the model (e.g., finding outputs the model might confuse for each other) [27]. Global explaining approaches take the model itself into account. An example would be the calculation of feature importances in Random Forests [28]. Model-independent approaches can be applied on every classifier, while model-specific explaining algorithms are designed for one type of classifiers, e.g., SVMs or CNNs. Two examples for local explaining algorithms are Sensitivity Analysis (SA) and Layer-Wise Relevance Propagation (LRP). They both produce a heatmap with pixels relevant to the classification [29]. This study compares and evaluates different CNN architectures upon their ability to correctly identify cardiomyocyte’s differentiation status/quality. Furthermore, a comparison between a 2D and 3D analysis of fluorescence images is made and explainability methods will be applied onto the classifiers to investigate the reasoning for a certain cellular stratification. In the end, it will be examined whether the predictions match the biological criteria for differentiated cardiomyocytes.

## Results

### Individual model development and classification

Figure 1 shows the categorical accuracies, validation accuracies, and confusion matrices on the test set for the singular 2D model on all classification tasks. All accuracies have a rather logarithmic progression over time, a typical training curve. For sarcomerisation classification, the training accuracy reached a plateau at almost 100 % after around 100 epochs. It took the validation accuracy around 80 epochs to reach a value of around 70 %, where it remained for the rest of the training epochs. The confusion matrix for sarcomerisation classification has its highest values along the diagonal, with the next highest values in row 2, column 1 and 3, which means that most of the data set is correctly classified. The accuracy of sarcomerisation classification on the test data set is 68.68 %. The training accuracy for directionality classification settled at almost 100 % after around 150 epochs. The validation accuracy reached a value of around 60 % after approximately 120 epochs. Again, the highest value per row lies along the diagonal of the confusion matrix. Directionality classification has an accuracy of 57.95 % on the test data set. Cell source classification’s training accuracy reached a plateau of almost 100 % after about 110 epochs. The validation accuracy settled at around 75 % after approximately 130 epochs. In the confusion matrix computed on the test data set, the highest values are located on the diagonal. The accuracy of cell source classification on the test data set is 74.85 %.

**Figure 1.**
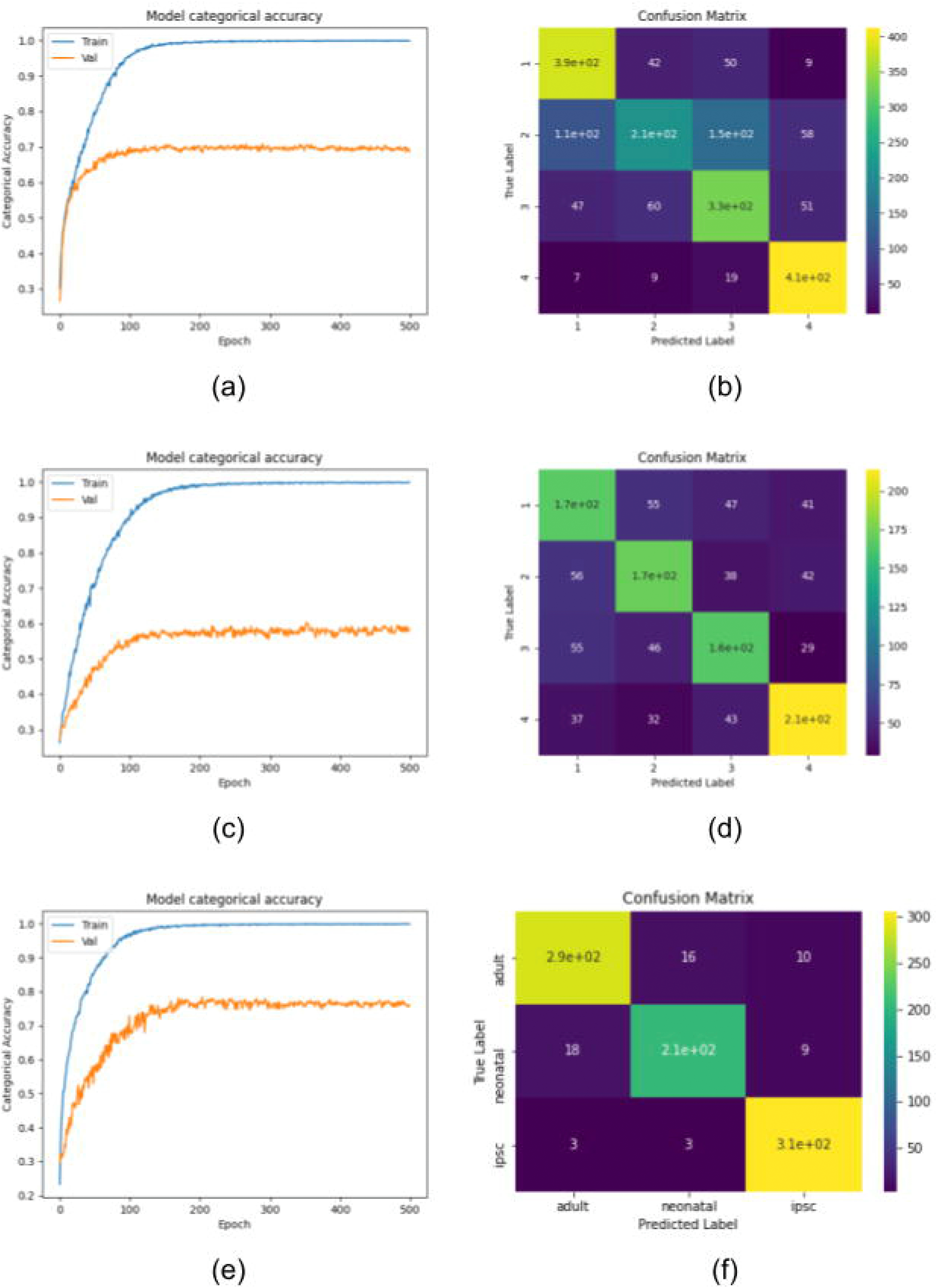
Accuracy plots and confusion matrices for the singular model. Shown are the categorical training (train, blue) and validation (val, orange) accuracies over time as well as the confusion matrices computed on the test set. (a-b): Sarcomerisation classification; (c-d): Directionality classification; (e-f): Cell source classification

In Fig. 2, the confusion matrices computed on the test set for the 2D ensemble model are shown. For all three classification tasks, the highest value per row lies on the diagonal. Sarcomerisation and directionality classification each have one cell classified as none of the ratings. The accuracy of sarcomerisation classification on the test data set is 70.93 %. It is 64.16 % for directionality classification and 82.06 % percent for cell source classification. As the individual classifiers were trained successively, no single training/validation accuracy for the whole model can be computed. They are, however, computed for each individual classifier in the model. The first sarcomerisation classifier, a one-vs-all model for rating ”1”, has a final training accuracy of 99.78 % and a final validation accuracy of 91.10 % after 1000 epochs of training. The second classifier, for rating ”2” respectively, settled at a training accuracy of 99.53 % and a validation accuracy of 85.23 %. For classifiers 3 and 4, the final training accuracies are 97.12 % and 99.56 %, while the final validation accuracies are 78.44 % and 92.90 %, respectively.

**Figure 2.**
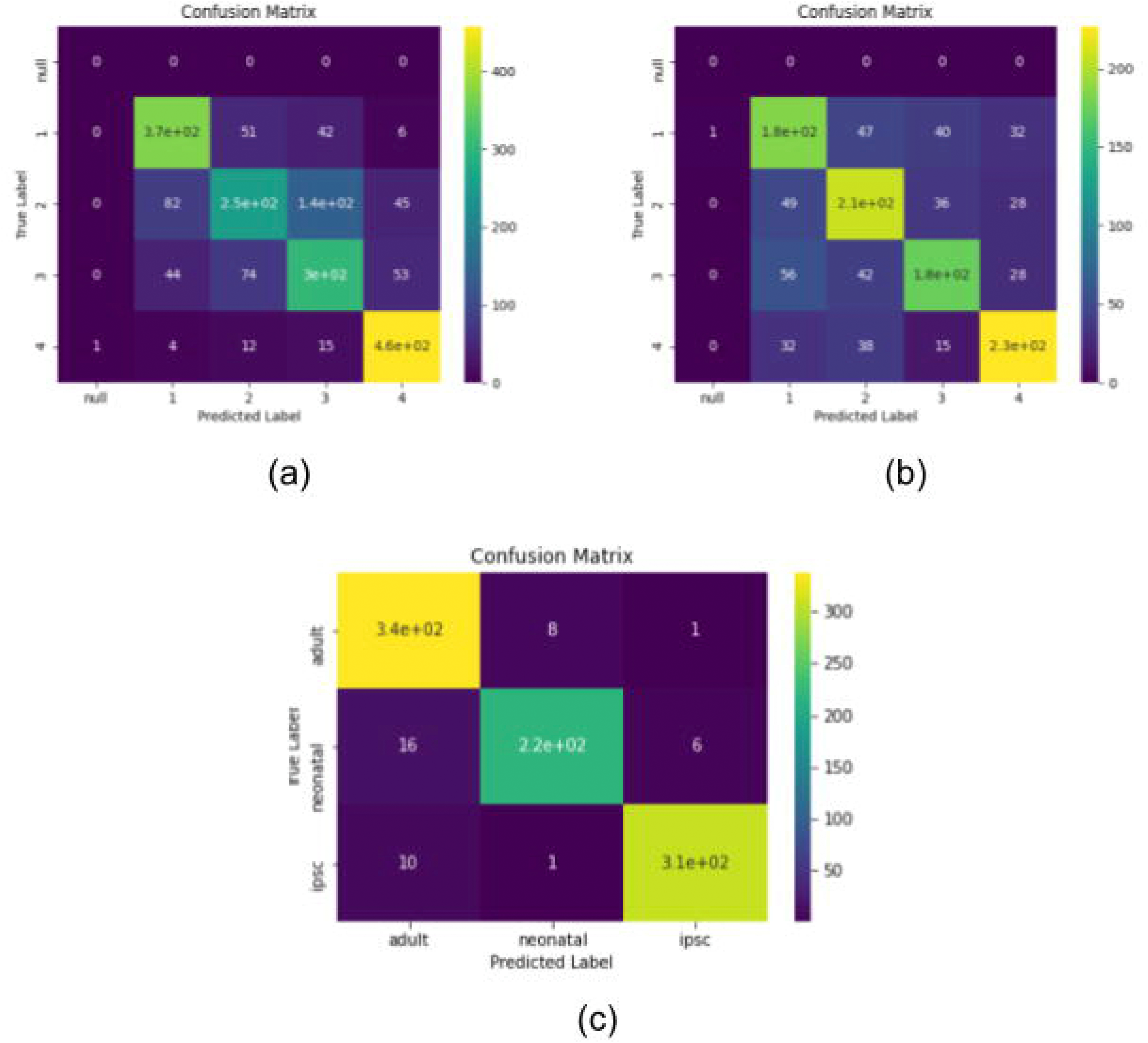
Confusion matrices for the one-vs-all ensemble model. Shown are the confusion matrices computed on the test set. (a): Sarcomerisation classification; (b): Directionality classification; (c): Cell source classification

The first directionality classifier has a final training accuracy of 99.66 % and a final validation accuracy of 84.44 %, each after 1000 epochs of training. For classifiers 2 and 3, final accuracies settled at 99.75 % and 99.16 % (training) and 81.91 % and 81.33 % (validation). The fourth directionality classifier reached a final training accuracy of 96.89 % and a final validation accuracy of 84.79 %.

The first cell source classifier is an one-vs-all model for the cell lineage ”adult”. It has a final training accuracy of 99.96 % and a final validation accuracy of 97.10 % after 1000 epochs of training. For the ”iPSC” classifier, the final training accuracy is 99.89 % and the final validation accuracy 97.53 %. The training accuracy for the ”neonatal” classifier reached 99.67 %, while the validation accuracy reached 92.84 %, each after 1000 epochs.

The accuracy plots for the 3D model based upon the singular model, also follow a typical training trend (see Fig. 3). For sarcomerisation classification, the training accuracy reached 100 % after 50 epochs, while the validation accuracy remained at around 78 % after 100 epochs. The confusion matrix has its highest values along the diagonal, which means that the vast majority of images (90.91 %) were correctly classified. For directionality classification, the training accuracy is almost 100 % after 500 epochs. The validation accuracy reaches a peak at about 68 % after around 150 epochs and then slowly declines to around 62 % after 500 epochs. The accuracy of directionality classification on the test data set is 62.60 % and the confusion matrix has its highest values along the diagonal with a notable peak in the last element of the second row. The training accuracy for cell source classification is 100 % after 500 epochs of training, while the validation accuracy settles at around 80 % after 70 epochs. In the confusion matrix computed on the test set, the highest values are located along the diagonal and in the second column of rows 4 and 5. The accuracy for cell source classification on the test set is 79.04 %.

**Figure 3.**
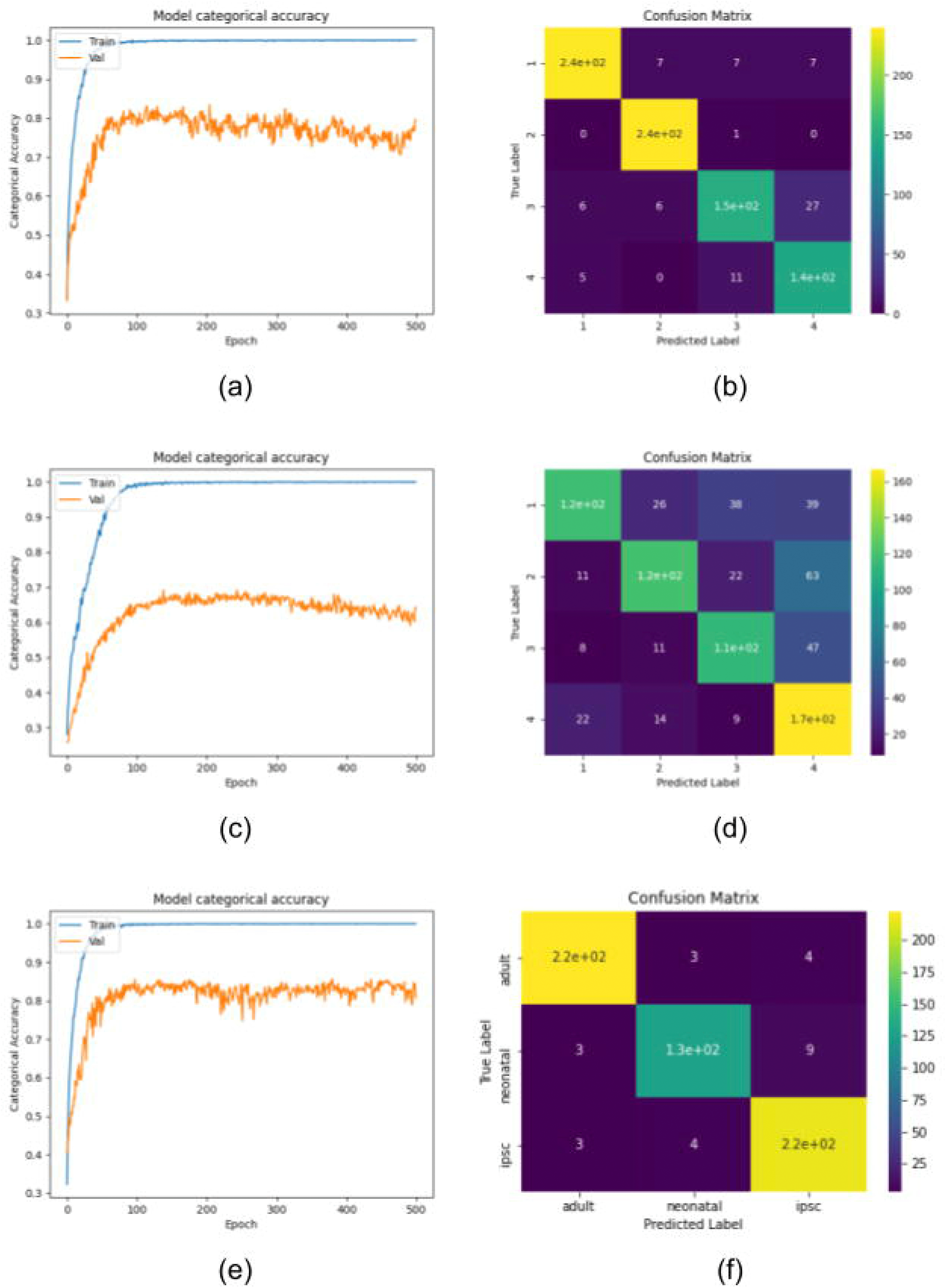
Accuracy plots and confusion matrices for the 3D model. Shown are the categorical training (train, blue) and validation (val, orange) accuracies over time as well as the confusion matrices computed on the test set. (a-b): Sarcomerisation classification; (c-d): Directionality classification; (e-f): Cell source classification

### Performance Comparison

The weighted f1-scores for the classifiers evaluated in this study are summarized in Fig. 4. It is different from the normal multi-class f1-score, as the per-class f1-scores are not only averaged, but also weighted according to the occurrence of their respective class. For the singular 2D model, directionality classification performs with a weighted f1-score of 57.96 %, while sarcomerisation classification reaches 67.33 %, and cell source classification reaches 74.81 %. Sarcomerisation classification yields the second best results for the 2D ensemble model (70.16 %), while directionality classification reaches 64.19 %. This model performs best on cell source classification by achieving a weighted f1-score of 82.06 %. Directionality classification for the singular 3D model has a weighted f1-score of 62.63 %, while the weighted f1-score for cell source classification is 79.14 %. The weighted f1-score for sarcomerisation classification lies at 90.87 %. Table 1 shows the test accuracies and weighted f1-scores computed on the test data set for all evaluated models.

**Table 1.** Test accuracies and weighted f1-scores for the models evaluated in this study. All values are presented in percent [%] and were computed on the respective test set. For the two transfer models, only the final results after retraining all layers are presented. In each column, the lowest and highest values are highlighted. (sarc: sarcomerisation, dir: directionality, cs: cell source)

|  | Test accuracy |  |  | Weighted f1-score |  |  |
| --- | --- | --- | --- | --- | --- | --- |
|  | sarc | dir | cs | sarc | dir | cs |
| 2D singular model | 68.68 | 57.95 | 74.85 | 67.33 | 57.96 | 74.81 |
| 2D ensemble model | 70.93 | **64.16** | **82.06** | 70.16 | **64.19** | **81.94** |
| 3D singular model | **90.91** | 62.60 | 79.04 | **90.87** | 62.63 | 79.14 |

**Figure 4.**
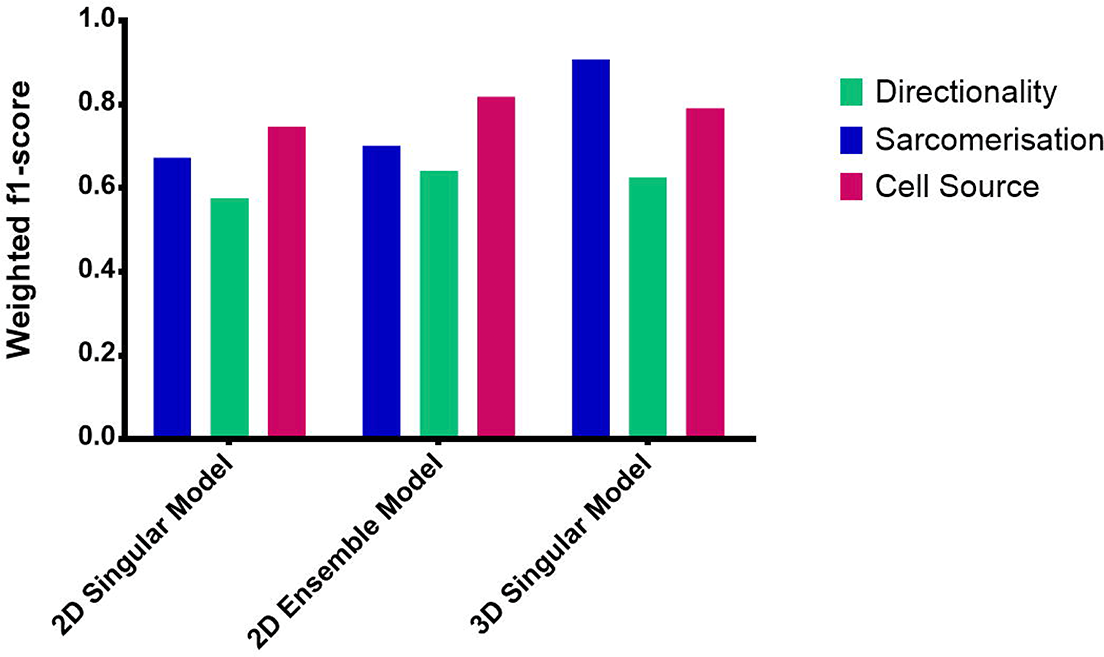
Comparison of the weighted f1-scores between all evaluated models.

### Explainability Heatmaps

In order to understand the reasoning behind the singular 2D models decisions, the explainability algorithms iGrad and Grad-CAM were applied. In Fig. 5a, heatmaps produced by IGrad as well as Grad-CAM for the singular 2D model can be seen. It correctly classified these images with a rating of ”4” for sarcomerisation, meaning this particular cell has a high degree of sarcomerisation. The heatmaps produced by IGrad have generally lower values than those produced by Grad-CAM.They lack areas of high importance, as the cells themselves have no impact on classification. There are however, slight accumulations around the edges of cells. When looking at the Grad-CAM heatmaps, the highest values lie inside the cell, with peaks around the edges. Almost the whole cell has a positive impact on classification. The heatmaps for the other classes can be seen in the supplemental material.

**Figure 5.**
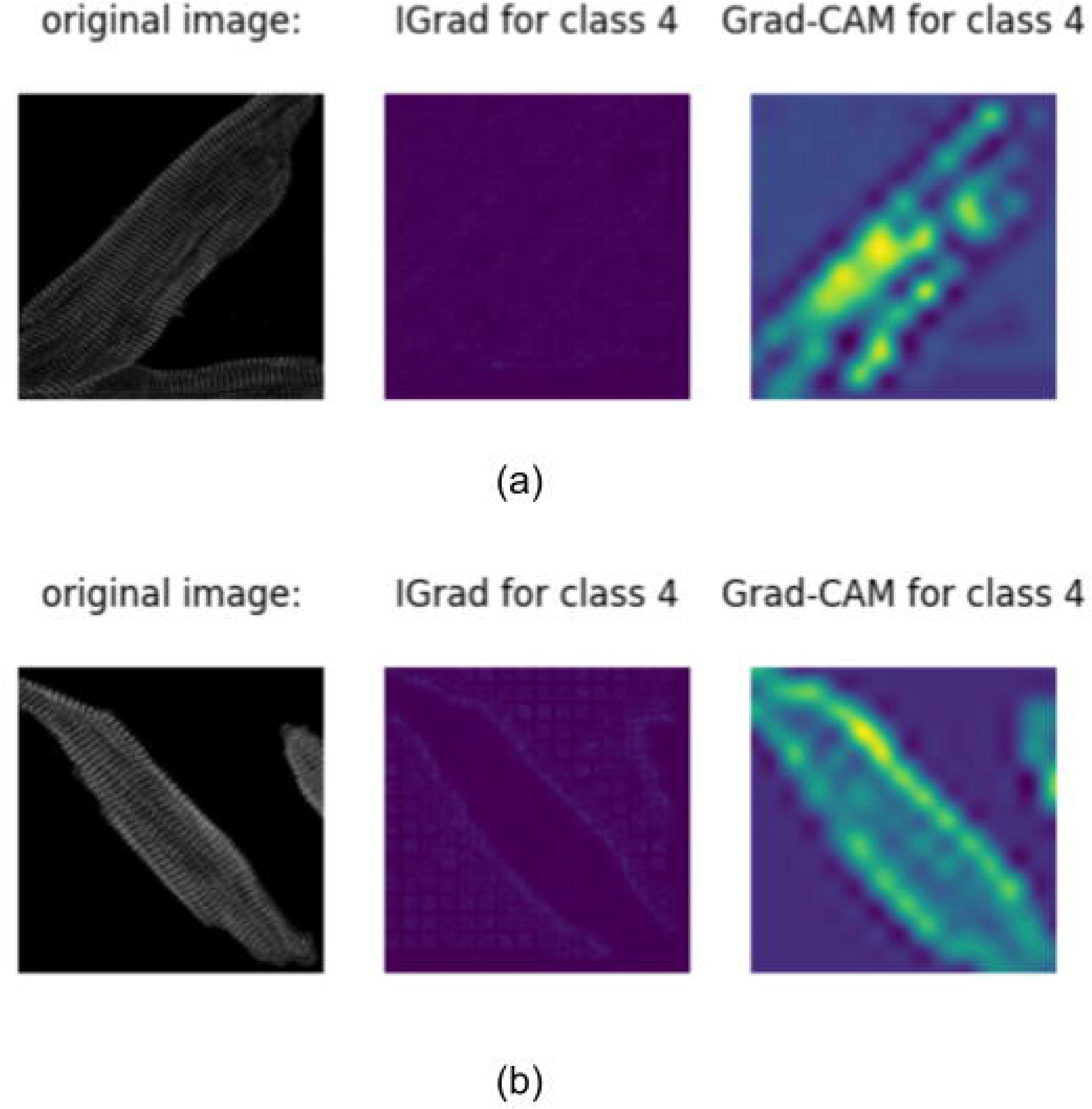
IGrad and Grad-CAM heatmaps for a) sarcomerisation and b) directionality rating ”4”. Images were correctly classified as ”4” by the singular 2D model.

The heatmaps Grad-CAM produced for the directionality have their peaks located inside the cardiomyocytes, while the one produced by IGrad barely have peaks at all (Fig. 5b). The IGrad heatmaps have very few values greater than zero.

The heatmaps produced by IGrad for the cell source have only very few values greater than zero. Inside the cells, there are none visible. The Grad-CAM heatmaps have their highest values inside the cells for classes ”adult” and ”iPSC” and on the cellular border for class ”neonatal” (see Fig. 6). Note that for ”adult” and ”iPSC”, Grad-CAM also highlights the borders of a cell.

**Figure 6.**
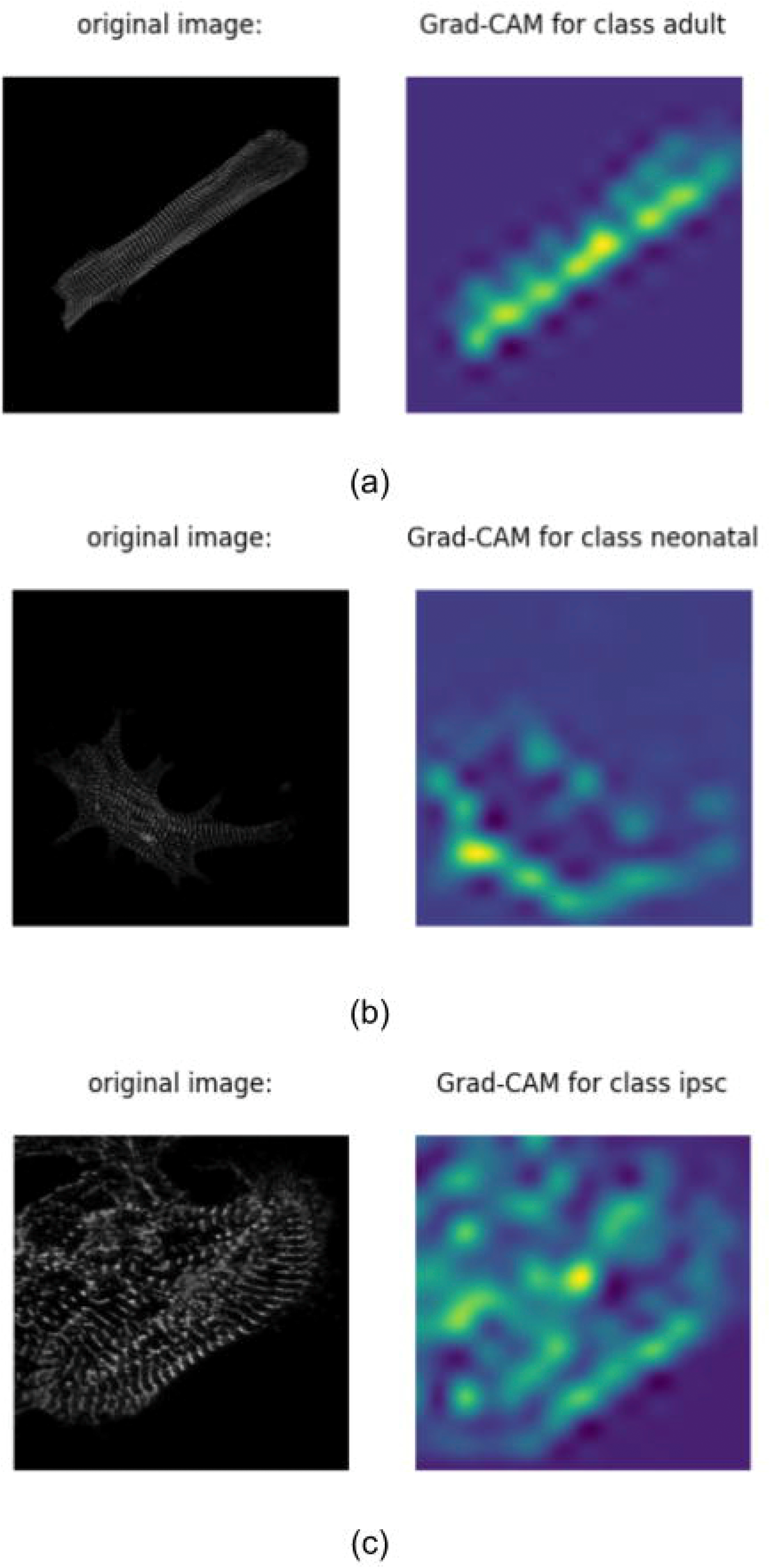
Grad-CAM heatmaps for cell source classification. Images were correctly classified by the singular 2D model.

## Discussion

### Cardiomyocyte quality is determinable via CNNs

When looking at the accuracies and the f1-scores of the models, it is clear that in principle, the evaluation of cardiomyocytes using DL is possible. This is a promising result, as in past approaches, neural networks mainly delivered well-suited results on bright field microscopy images, as opposed to fluorescence images [20, 25]. The classification regarding sarcomerisation, directionality, and even cell source yields encouraging findings in terms of accuracy. Because of the balanced data set created by data augmentation, the baseline classification accuracy (the model predicts every data point to belong to the majority class) lies at around 20 % for sarcomerisation and directionality classification and at around 30 % for cell source classification. For all tasks, these baseline accuracies can be considerably outperformed. It also appears that a shallow model is sufficient to tackle all classification tasks. The singular 2D model has far less parameters than other, established image classification networks like the VGG-16 or MobileNet [30]. This suggests that the features extracted by a CNN do not need to be overly specific or detailed to ensure an accurate classification of either sarcomerisation, directionality, or cell source. From a human domain expert perspective, the rating of sarcomerisation and directionality are intuitively easy, as humans can easily spot parallel patterns and fractions of objects. Both are relatively easy features and so it is quite surprising that directionality classification consistently performs worse than sarcomerisation and cell source classification. However, directionality classification, when looked at in detail, are two separate tasks. First, the main cellular axis has to be found, which may be easy for adult cardiomyocytes, but has its challenges for the other analyzed cell sources. Second, the sarcomere structures must be evaluated according to their axis, potentially leading to consequential errors, if the main axis is not determined properly. This may be an explanation to why it is more difficult for neural networks to evaluate on directionality than on sarcomerisation or cell source. It has been shown that cardiomyocytes generated from iPSCs resemble fetal cardiomyocytes rather than adult ones [31]. Our analysis also confirms this finding, as all three analyzed cell types can be distinguished from each other, meaning that there is still an observable difference between primary and generated cardiomyocytes(see Figs. 1–3).

Stochastic Gradient Descent, although being a widespread optimizer, has its problems with large data sets and/or high dimensional feature space [32, 33]. The latter, high dimensional feature space, holds true for the data used in this work, especially for 3D classification. The Adam optimizer tackles these problems and was consequently used for all models. The learning rate was set to 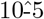 and made adaptive. Adaptive learning rates have been shown to boost classification results [34], which was also the case here. Another approach, increasing the batch size over training instead of decreasing the learning rate, was not applied, although it showed promising results in previous works [35]. For the applications in this study, it was not applicable, as the batch sizes cannot be increased arbitrarily because the images are too large in file size, especially for the 3D classification. Although transfer models have not been evaluated in this work, they have been applied onto cellular image analysis tasks in the past. For example, Dong et al. were able to correctly distinguish malaria infected cells from healthy ones using bright field microscopy with an accuracy of up to 98.13 % [20]. Cascio et al. built a transfer model, which was able to classify indirect immunofluorescence images of Human Epithelial type 2 (HEp-2) into fluorescence intensity classes with an accuracy of 93.80 % [36]. HEp-2 is a marker for antinuclear antibodies and, therefore, autoimmune diseases. Both approaches made use of the AlexNet [37], which is rather simple in terms of architecture and number of parameters.

### 3D classification outperforms 2D classification in terms of accuracy

The comparison between the 2D singular model and the 3D model, two similar models with different dimensionalities, shows a slight improvement in the classification accuracies when 3D images were analysed. An increase in model performance comes to no surprise, as the images are bigger and able to store more information in the additional dimension. However, they do not seem to become more complicated, as the same shallow architecture is able to extract features, which suffice for classification. The addition of a third dimension improves the classification marginally for directionality and cell source classification (about 5 % each) and drastically for sarcomerisation classification (about 20 %). The computation time however increases drastically. Training a classifier on 2D data for 500 epochs is a matter of hours, on 3D data, it takes days. Interestingly, both 2D and 3D models take up about the same storage space with around 6 megabytes. Still, the increase in computation time, not only for the training of the classifier, but also for image preprocessing, outweighs the improvement of classification accuracy at this point. There are two options for the application of this evaluation: Use the 2D classifier, and get quick, but slightly imprecise results, or use the 3D classifier, and get more sensitive and accurate results at the expense of computation time. As computational hardware is constantly improving, the difference in computation time might be significantly reduced in the future. 3D analysis has been long proven to outperform 2D analysis on a wide range of tasks[38]. In the life sciences, 3D image analysis with DL is often used for segmentation [39, 40, 41]. In this context, it has been shown that 3D segmentation outperforms the segmentation of all individual 2D slices [42]. 3D image classification approaches for biological images are rare. One example is the classification of functional connectomes by Khosla et al. They were able to correctly classify 73.30 % of functional magnetic resonance images (fMRI) using a custom build model trained on the Autism Brain Imaging Data Exchange (ABIDE) data set [43]. This is similar to the accuracies reached by 3D classification in this study (62.60 % to 90.91 %).

### Cellular borders are of high interest

The implementation of IGrad leads to no interpretable results. This could be due to an error in the implementation of IGrad in the keras-explain package. Other possible error sources could be the shallowness of the evaluated model, the proximity of a monochrome cellular image to the black image baseline or a combination of these two. In contrast, the heatmaps produced by Grad-CAM are very informative, although they lack detailed spatial information of pixel-wise impact on the classification. This is due to the shallow architecture of the evaluated model. The deeper a CNN is, the more specific the features in the last feature map are. The singular model simply seems not deep enough to extract detailed features from the images. Still, it is apparent that the most important regions for all classification tasks are within the cell. This is not surprising, as the sarcomere network lies within the cell and thus, all valid decisions based upon this sarcomere network must be traceable to the cell itself. Interestingly, there are outlier images, though, whose Grad-CAM heatmaps locate the important regions to be outside of the cell. This could be due to ”zero-filters”, convolutional filters that learn to find background or, as in this case, black regions in the images. Especially for sarcomerisation, these filters can be useful, as they allow for reverse explanations. If a large part of the image and/or the cell is black, the sarcomerisation rating will probably be low. Cellular borders seem to be of high importance according to the heatmaps produced by Grad-CAM. For all three classification tasks one can find examples of this (e.g., Fig. 5, Fig. 6). This could hint at edge detection being learned by the networks. For sarcomerisation classification, this makes sense, as the network may recognize where the cells are on the image and, consequently, which areas in the image should have a dense sarcomere network. Adult cardiomyocytes tend to be elongated and thin, while cardiomyocytes generated from iPSCs resemble fetal cardiac muscle cells, which are more likely to be round or have irregular shapes [44]. Therefore, edge detection could benefit the distinction between these cell sources. In general, Grad-CAM produced heatmaps of the singular model that seemed to highlight regions in the image that would also be deemed important by human curators.

## Conclusion

In this work, a novel unbiased tool to evaluate the sarcomerisation, directionality, and cell origin of a cardiomyocyte is presented. Several different CNNs were trained on 2D and 3D fluorescence images of cardiomyocytes with the sarcomere network stained. The cardiomyocytes used in this study were rated based on their sarcomerisation and the orientation of sarcomere structures (directionality) beforehand (CCRS scheme). The trained models were subsequently evaluated by feeding them into two explainability algorithms, IGrad and Grad-CAM, which highlight the areas in an image that are most important for the respective classification.

IGrad andGrad-CAM both produce heatmaps, where IGrad did not provide interpretable results in our data. The heatmaps produced with Grad-CAM have their highest values inside the cell and at cellular borders for all three classification tasks, meaning that these regions are important for classification. However, these heatmaps do not contribute to novel findings, but highlight that cells are being recognized by the classifier.

In general, it is shown that cellular fluorescence images can be analysed with CNNs. A classifier was built that is capable of predicting 82 % of cardiomyocyte origins, 71 % of sarcomerisation ratings, and 64 % of directionality ratings correctly. This classifier can be used to make independant and trustworthy predictions on the quality of generated cardiomyocytes based on the sarcomere network. This underlying work will significantly benefit the unbiased evaluation of cardiomyocytes, as a fast and reliable tool for cardiomyocyte aggregates is now available.

## Methods

In this study, 2D and 3D fluorescence images of cardiomyocytes were analyzed with CNNs. These CNNs were trained to distinguish between different ratings for cardiac muscle states, which were assigned to the cells beforehand. The CNNs were evaluated according to their accuracy and f1-score. Additionally, the best performing model was analyzed with explainability algorithms in order to visualize the network’s behaviour and criteria for classification. The workflow is shown schematically in Fig. 7.

**Figure 7.**
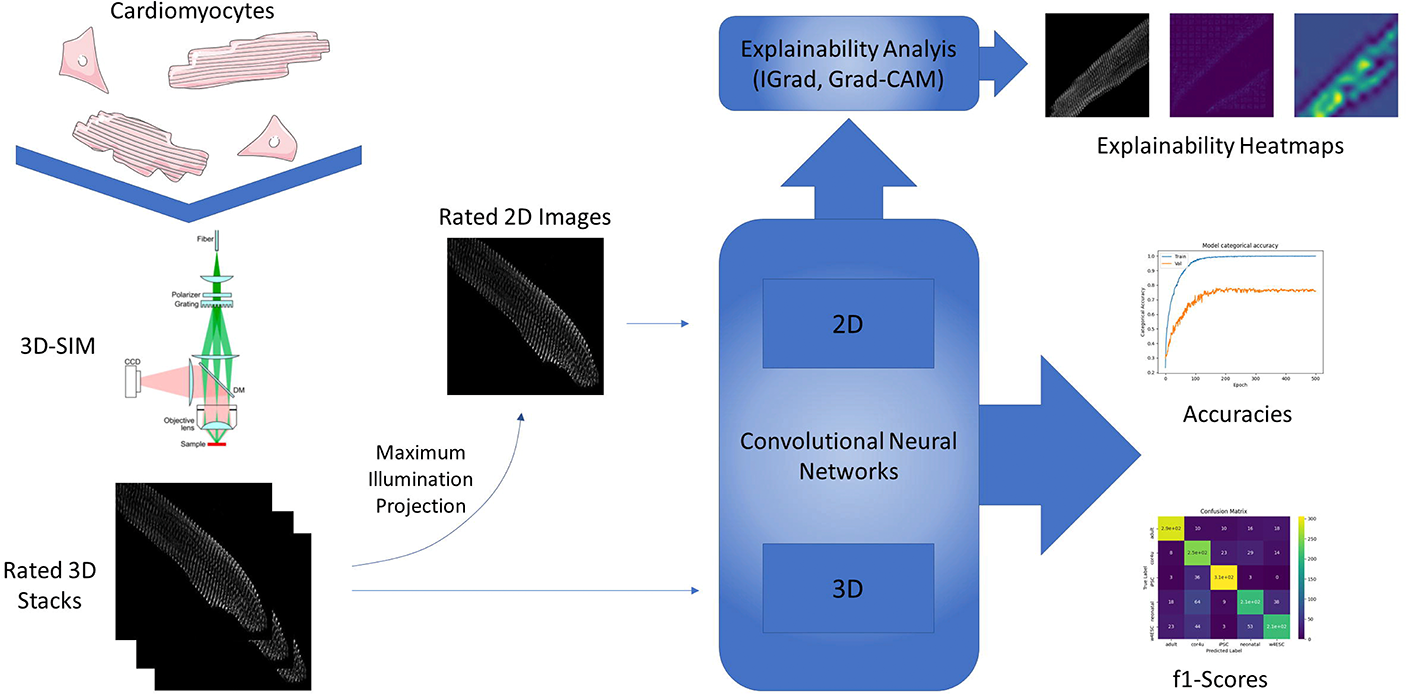
Schematic overview of the study design. SIM: Structured Illumination Microscopy, IGrad: Integrated Gradients,Grad-CAM: Gradient-Weighted Class Activation Mapping

### Cell origin and image acquisition

A comparison of murine-derived cardiomyocytes (adult and neonatal) and human-derived induced pluripotent stem cells (iPSCs), which both differentiated into cardiomyocytes, serve as the basic sarcomere models in this work. This allows for a validation of the iPSCs, which were differentiated and harvested by the RTC.

The cellular images used in this study were acquired using 3D fluorescence structured illumination microscopy (SIM), which provides for resolutions of about 100 nm [45, 46]. Between 36 and 83 images have been taken and arranged into a z-stack per cell. The x- and y-sizes of the images range from 564 to 2002 pixels and from 392 to 2027, respectively, depending on the size of the cell captured. In order to obtain the 3D fluorescence images, the cells were stained as follows. At first, the cells were fixed by adding pre-warmed 4 % paraformaldehyde (PFA) directly into the culture medium with a ratio of 1:1 for five minutes at 37 °C. The cells were washed two times with phosphate buffered saline (PBS) for 5 min each. After this, they were permeabilized with 0.2 % Triton for five minutes and again washed twice with PBS for five minutes each. Next, the immunostaining took place. Therefore, the unspecific binding sites of the cells were blocked with 1 % bovine serum albumin (BSA) at room temperature for 60 min. The cells were then stained with the primary antibody against *alpha*-actinin (see Table 2), which is diluted in 1 % BSA, for 60 min at room temperature. *alpha*-actinin binds to the actin filaments of the sarcomeres and stabilizes the muscle contractile apparatus [47]. Two washes with 0.2 % BSA for five minutes each followed. Then, the cells were incubated with the secondary antibody (see Table 2), which was diluted in 1 % BSA, for 45 minutes at room temperature. Again, the cells were washed, twice with 0.2 % BSA and twice with PBS, each for five minutes. Coverslips were rinsed with distilled water and the cells were embedded on slides using mounting medium containing DAPI.

**Table 2.** Antibodies used for immunostaining of cardiomyocytes.

| Type | Antibody |
| --- | --- |
| Primary | Sarcomeric *alpha*-actinin antibody ea53<br>Abcam plc, USA |
| Secondary | F(ab')<sub>2</sub>-Goat anti-Mouse IgG (H+L) Cross-Adsorbed Secondary Antibody<br>Thermo Fisher Scientific, USA |

### Introduction of a cardiomyocyte cell rating system (CCRS)

The cells in the fluorescence images have been rated regarding the orientation of sarcomere structures relative to the longitudinal axis of the cell (directionality) and their sarcomerisation, which both correlate to the maturity of a cardiomyocyte. ”1” marks the lowest rating and ”4” the highest, with ”2” and ”3” as intermediate steps. A cell with sarcomere structures parallel to the cell’s longitudinal axis would be marked as ”1” for directionality, whereas a cell with perpendicular orientated sarcomere structures would be marked as ”4”. Similarly, cells with a high degree of sarcomerisation were marked as ”4”, as opposed to ones with a low level, which were marked as ”1”. Example images for all ratings can be seen in Fig. 8.

**Figure 8.**
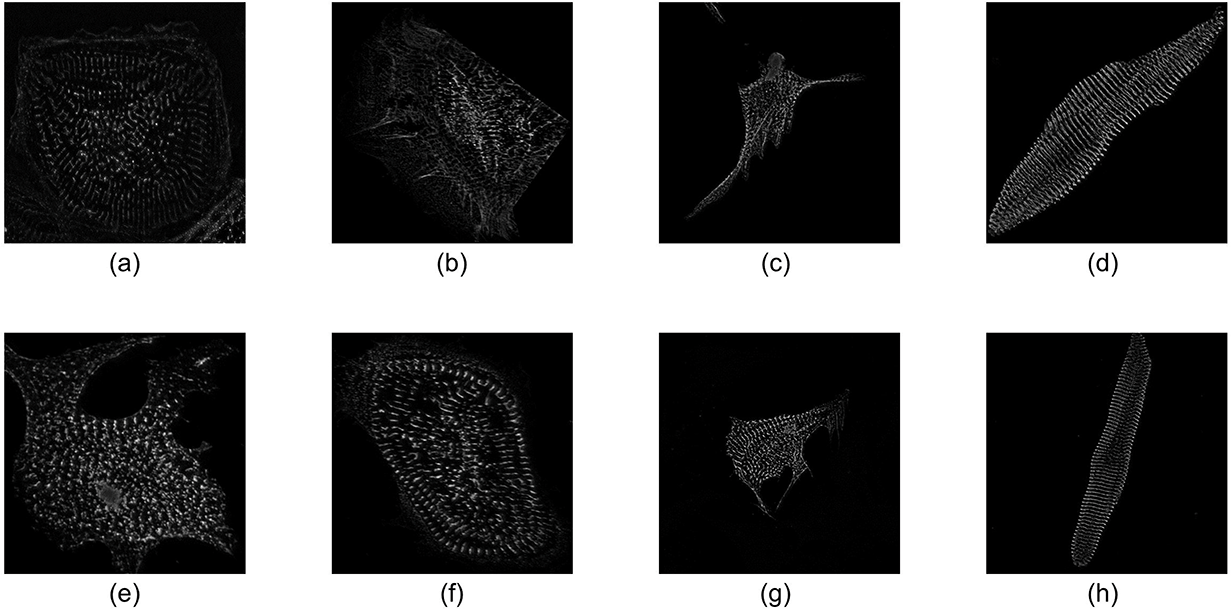
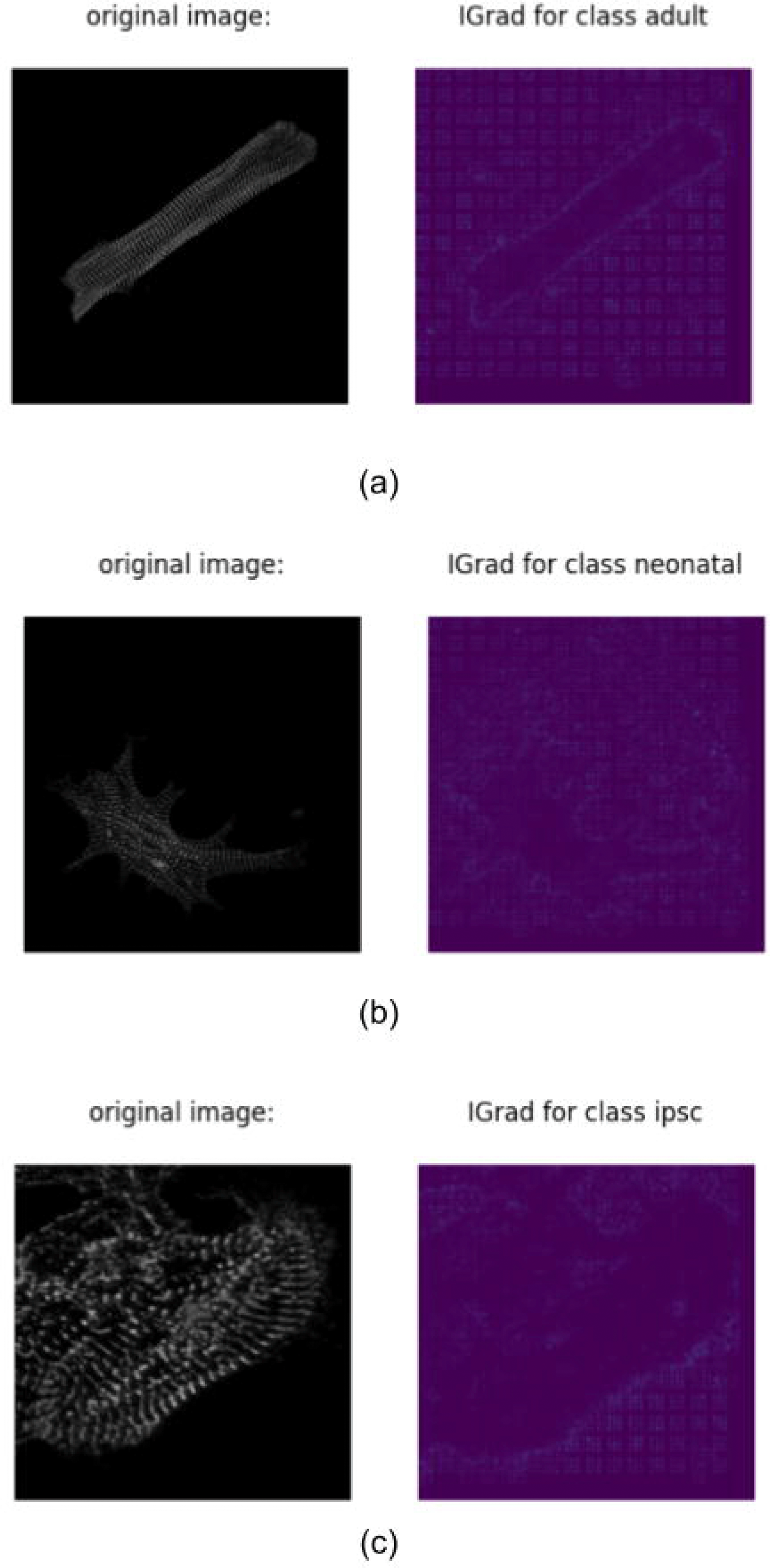
Example images for all different sarcomerisation and directionality ratings.

### Cardiomyocyte image processing, classification, and explaination

As the original images are three-dimensional, 2D images had to be created with ImageJ (version 1.52a) [48]]. The common practice of reducing the dimension by a maximum illumination projection along the z-axis was used to obtain two-dimensional images of the cardiomyocytes fluorescence stacks [49]. Image preprocessing consisted of resizing the images to a models respective input size and scaling the pixels between 0 and 255. Data augmentation was implemented to increase the number of data points. The augmentation was performed depending on the relative occurrence of a class to simultaneously guarantee for a balanced data set. The class with the fewest data points was multiplied by a factor of 15 for 2D classification and 9 for 3D classification. This is because more augmented 3D images exceed the available storage space. Images were either flipped horizontally, vertically, rotated by a random degree or a combination of these three methods. This does not distort the image and allows validation of the results [50].

Two different 2D classification approaches will be presented in the following: one singular model with varying architectures and one ensemble model. The first model is shallow, and consists of only three convolutional blocks, and, thus, has only a few parameters. The second model makes use of an ensemble of neural networks. For each class, a binary classifier was trained to distinguish between this class and all other classes.

3D image classification is often accompanied by a task like depth perception (e.g., human pose estimation [51]) or shape reconstruction (mesh/point cloud classification [52]). There are few cases, where 3D images are classified as a whole, mainly for medical purposes, but these approaches lack a common model used. Therefore, a native 3D classification model has been evaluated. The 3D model closely resembles the singular model. It is also made up from three blocks of 3D convolution, batch normalization, max pooling, and dropout layers. The detailed architecture can be seen in the appendix. The network was trained with the Adam optimizer with a learning rate of 10-5, which decays by 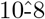 each epoch.

Table 3 shows a comparison between the parameters of each model. The number of parameters strongly influences the storage space of a model, as well as the time needed to train and test it. Models with fewer parameters are easier to implement and take less time to make predictions.

**Table 3.**
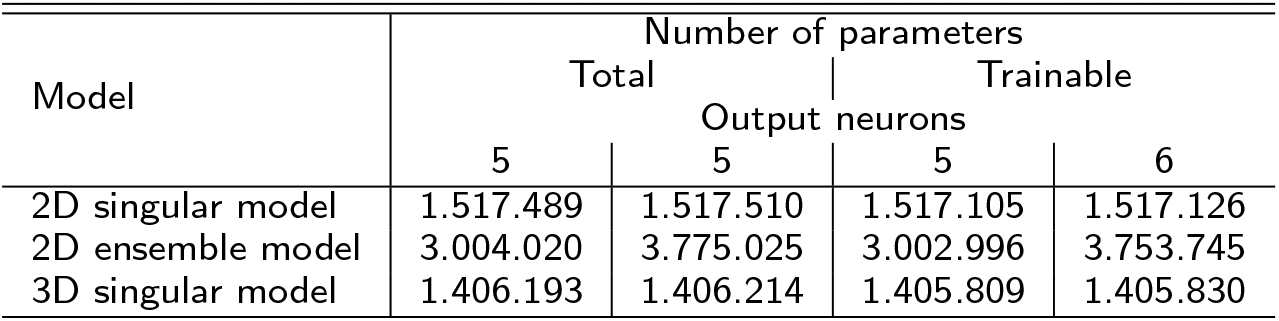
Comparison of the number of parameters between the different models used. For the ensemble model, the sum of the individual models has been calculated.

Two different methods for explainability analysis were applied in this thesis: Integrated Gradients (IGrad, [53]) and Gradient-weighted Class Activation Mapping (Grad-CAM, [54]). Both methods allow for a pixel-wise decomposition of CNNs. Here, we made use of the keras-explain 0.0.1 package for Python 3.7 to implement both methods and compute the heatmaps [55]. The heatmaps were scaled between 0 and 255.

All models were initialized with random weights and biases following a uniform distribution around zero. Native models were each trained for 500 epochs. The code was implemented in Python 3.7.2 using Keras 2.1.6 with Tensorflow backend and scikit-learn (version 0.20.3) packages [56, 57, 58]. Tests were run on an nVidia GeForce GTX 2080 GPU with 8 GB RAM.

## Ethics approval and consent to participate

Not applicable.

## Consent for publication

Not applicable.

## Availability of data and materials

The datasets used and analysed during the current study are available from the corresponding author on reasonable The python code used for analysis is available at github [59].

## Author’s contributions

Conceptualization, MH, TM, MW and OW; Formal analysis, MH; Funding acquisition, OW and RD; Investigation, MH and HL; Methodology, MH, HL, RD and MW; Supervision, RD, TM and OW; Visualization, MH; Writing — original draft, MH, HL and MW

All authors have read and agreed to the published version of the manuscript.

## Acknowledgements

Not applicable.

## Additional Files

Additional file 1 — Singular2D.png

Architecture of the singular 2D model in Keras notation.

Additional file 2 — Ensemble2D.png

Architecture of the ensemble 2D model in Keras notation.

Additional file 3 — Singular3D.png

Architecture of the singular 3D model in Keras notation.

Additional file 4 — CellSource-IGrad.png

IGrad heatmaps for cell source classification. Images were correctly classified by the singular 2D model.

